# A generative reference grammar of healthy TCR repertoires reveals cancer-associated immune remodeling

**DOI:** 10.64898/2026.04.29.721631

**Authors:** Archana Balan, Yuval Elhanati, Kevin E. Meza Landeros, Marcus De Almeida Mendes, Jiaying Lai, Syed Shujaat Ali Zaidi, Mesut Unal, Betty Y.S. Kim, Calixto-Hope G. Lucas, Emerenziana Runco, Vinay K. Puduvalli, Jennifer Gantchev, Charles A. Whittaker, Pratibha Sharma, Viviane Tabar, Michael J. Cima, Gerard Baquer, David A. Reardon, Alexei Stortchevoi, Adrienne Boire, Linghua Wang, Forest M. White, Dimitrios N. Sidiropoulos, Kenny Kwok Hei Yu, E. Antonio Chiocca, Valsamo Anagnostou, Data Science TeamLab, Accelerating GBM Therapies TeamLab, Rachel Karchin

**Author notes:** To whom correspondence should be addressed: Rachel Karchin.

## Abstract

T-cell receptor (TCR) repertoires record how adaptive immunity is organized and how cancer and therapy reshape it, but this signal is hard to read: treatment-associated change is entangled with the V(D)J recombination constraints that shape every repertoire. We present CRAFT (Cancer Repertoire Anomaly Finding Transformer), a conditional sequence-to-sequence transformer that learns a nucleotide-level generative grammar of productive TCR-beta CDR3 sequences from healthy donors, conditioned on germline V(D)J assignments. A dual-head decoder mirrors the independence of V-D and D-J recombination, and curriculum training produces embeddings that define a healthy-reference coordinate system in which cancer-associated change appears as structured, measurable deviation. In proof-of-concept applications to a neoadjuvant checkpoint-blockade cohort sampled longitudinally across blood, and to serial single-cell profiling of T-cell subsets during oncolytic immunotherapy, CRAFT geometric metrics capture response-associated remodeling, including shifts in repertoire organization over time. On antigen-labeled benchmarks, CRAFT organizes specificity classes coherently, recovering structure that reflects shared antigen recognition.

## Introduction

T-cell receptor repertoires provide a high-dimensional readout of the immunogenicity landscape, encoding the baseline organization of adaptive immunity and its history of antigen encounters. Patterns of clonal expansion, V(D)J usage, and CDR3 sequence composition describe how T cell clonal repertoires are generated, selected, and maintained, and have emerged as promising biomarkers of immune competence and response to immunotherapy.^1,2^ High-throughput sequencing now profiles repertoires at scale, yet biologically meaningful structure is hard to extract. Each repertoire spans millions of distinct clonotypes, and the patterns that reflect V(D)J recombination and thymic selection^3,4^ must be distinguished from those that arise when disease and therapy reshape the repertoire. The probabilistic rules governing recombination have been characterized through generative models of TCR formation, from inference of per-sequence generation probabilities^5^ to efficient analytic computation across large repertoires^5–8^. These frameworks establish that physiologic recombination and selection impose broad statistical constraints on germline gene usage and CDR3 composition. Approaches for quantifying repertoire-level divergence between individuals and disease states have also been developed,^9–11^ but measuring how cancer-associated repertoires depart from healthy baseline architecture at the latent sequence structure level has remained an open problem. Meanwhile, antigen exposure in cancer and under immunotherapy can imprint distinct sequence patterns. Recent strategies, including CDR3 motif-based clustering,^12^ deep learning frameworks for TCR classification,^13–15^ and high-dimensional TCR embeddings,^13,15,16^ support tasks such as motif discovery and antigen-binding prediction, but most operate on amino-acid sequences and optimize for epitope-level classification rather than repertoire-level change detection. A principled framework for measuring how cancer repertoires depart from healthy baseline architecture has been lacking. We introduce CRAFT (Cancer Repertoire Anomaly Finding Transformer), a generative transformer trained exclusively on TCR-beta repertoires from individuals without known malignancy. Conditioned on germline V(D)J assignments, CRAFT reconstructs CDR3-beta nucleotide sequences, learning the latent recombination grammar and baseline manifold of CDR3-beta diversity. By operating at the nucleotide level, the model captures constraints partially obscured after translation, including codon-level structure and junctional microarchitecture. The resulting embeddings provide a reference coordinate system in which cancer-associated repertoires are quantified as structured deviations from expectation, enabling measurement of immune perturbation, convergence, and remodeling across time, tissue, and cellular compartments in TCR repertoires derived from cancer patients (Figure 1d). A deliberate scope constraint is the restriction to TCR-beta chains; although antigen recognition depends on the paired alpha-beta heterodimer, the vast majority of large-scale repertoire studies sequence only the beta chain, making it important that repertoire-level analytic frameworks can operate effectively in this setting.

**Figure 1.**
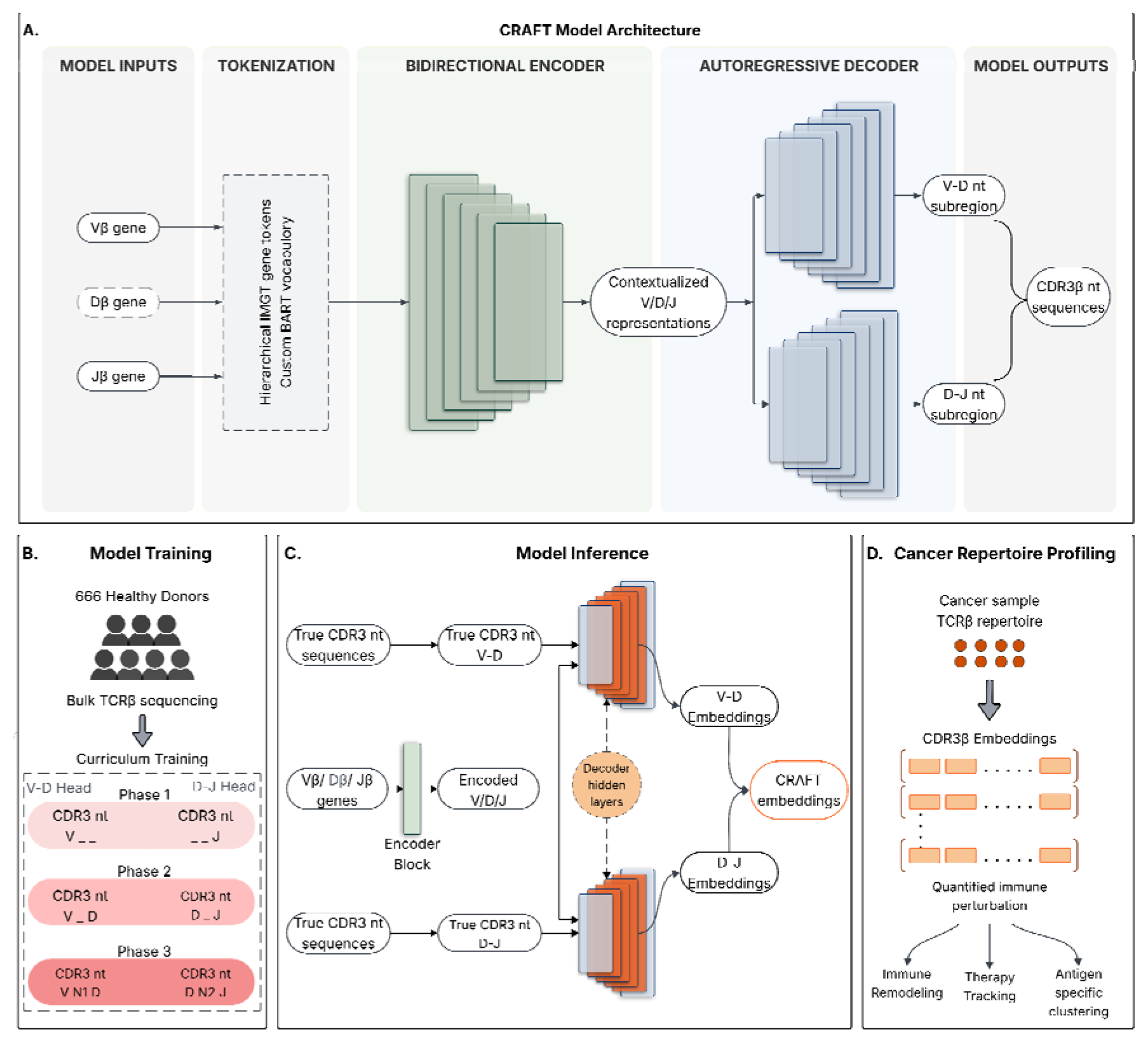
Overview of the CRAFT framework for learning a healthy-donor TCR embedding space and quantifying cancer-associated repertoire perturbation. (a) CRAFT model architecture. V, D, and J germline gene segments are tokenized using a hierarchical IMGT-based vocabulary (family, subfamily, allele) and encoded by a bidirectional encoder. An autoregressive decoder with two parallel output heads (V-D and D-J junctions) reconstructs CDR3-beta nucleotide sequences using character-level (A, C, G, T) tokenization, reflecting the known independence of V-D and D-J recombination events. (b) Model training on bulk TCR-beta repertoires from 666 healthy donors using multi-phase curriculum training with progressively increasing sequence complexity. (c) Model inference and embedding extraction via teacher-forced inference, with mean-pooled hidden states from both decoder heads concatenated to form the full CRAFT embedding. (d) Cancer repertoire profiling in the healthy-donor-derived reference space using geometric metrics.

## Design

Several architectural choices follow from this design goal. The model operates at the nucleotide rather than the amino-acid level. Translation to protein discards codon degeneracy, junctional diversity signatures, and palindromic (P) nucleotide patterns that carry information about the recombination process itself^5^; because separating recombination structure from selection-driven structure is the core aim, these signals are retained (STAR Methods). CRAFT adopts a BART-style conditional sequence-to-sequence architecture, coupling a bidirectional encoder over the germline V, D, and J segments with an autoregressive decoder that generates the non-templated CDR3-beta sequence (Figure 1a). We chose this conditional encoder-decoder over two common alternatives. A decoder-only transformer would require serializing the germline context into a single left-to-right token stream, whereas a bidirectional encoder that contextualizes all three segments before generation better reflects the fact that V, D, and J jointly constrain CDR3 structure; the distinct encoder and decoder vocabularies reinforce a design with separate input and output stages. A variational autoencoder was also considered but is less suitable here, because its embeddings are stochastic samples from an approximate posterior whereas our geometric metrics require deterministic, reproducible representations, and because its KL-divergence penalty would smooth precisely the fine-grained, germline-conditioned structure the framework is designed to detect (STAR Methods). Because V-D and D-J recombination events are largely independent, the decoder is split into a V-D head and a D-J head that share decoder hidden states but predict their junctional subregions independently, a design that improved convergence and reduced perplexity at the more variable D-J junction relative to a single-head variant. The encoder uses a hierarchical IMGT-based gene vocabulary in which each V, D, or J gene is decomposed into family, subfamily, and allele tokens (e.g., TRBV7-2*01 becomes [TRBV7, TRBV7-2, TRBV7-2*01]). This hierarchical encoding allows the model to share statistical strength across related genes while retaining allele-level resolution. The decoder uses a character-level nucleotide vocabulary (A, C, G, T) augmented with start, end, and padding tokens, operating directly on the DNA sequence rather than on amino acids. The model is trained with a multi-phase curriculum that introduces CDR3 length and gene-combination complexity progressively to stabilize learning. Sequence embeddings were obtained via teacher-forced inference^18,19^, in which the true V, D, J gene tokens and CDR3 subregion nucleotide tokens were provided to the model at each decoding step. Hidden states from the V-D and D-J decoder branches were mean-pooled across tokens and concatenated to form a unified per-TCR representation (Figure 1c). This approach extracts the model’s internal representation of each sequence given its full recombination context, yielding embeddings that encode how well each CDR3 conforms to the learned grammar. (STAR Methods).

Using the healthy reference embedding as a coordinate system, CRAFT supports several complementary analyses of cancer-associated repertoires (Figure 1d): (1) global repertoire remodeling, quantified through embedding variance, median distance of dominant clonotypes from their centroid, and longitudinal centroid shifts; (2) compartment-specific remodeling in single-cell data, quantified as enrichment of clonotypes displaced beyond a healthy-derived baseline threshold within defined T cell compartments; and (3) antigen-class organization, assessed by the separation of pathology-associated TCRs in the reference space. Each analysis measures a distinct geometric aspect of how a repertoire departs from healthy expectation (STAR Methods).

## Results

### CRAFT model training and evaluation

CRAFT was trained on bulk TCR-beta sequencing data from 666 healthy donors, comprising approximately 131 million productive rearrangements (Figure 1b). The cohort includes both CMV-seropositive and CMV-seronegative donors, as the Emerson et al. dataset^20^ was designed to capture the breadth of CMV-associated repertoire variation. We verified that this covariate does not dominate the embedding space (Figure S1), confirming that CRAFT embeddings reflect TCR sequence features rather than CMV serostatus. The dual-head decoder architecture improved convergence and better reflected recombination logic compared to a single-head variant, particularly for the more variable D-J junction, where the single-head model showed systematically higher perplexity (Figure 1a-c). On held-out sequences, CRAFT achieved low perplexity across germline-encoded regions, with uncertainty increasing in N-insertion regions consistent with their greater sequence variability (Figure 2a), indicating that the model distinguishes templated from non-templated sequence elements. Generated CDR3 length distributions closely matched empirical distributions within each VJ stratum, and nucleotide k-mer composition showed low Jensen-Shannon divergence across all orders tested (Figure 2b), demonstrating that CRAFT reproduces both global length constraints and local sequence composition.

**Figure 2.**
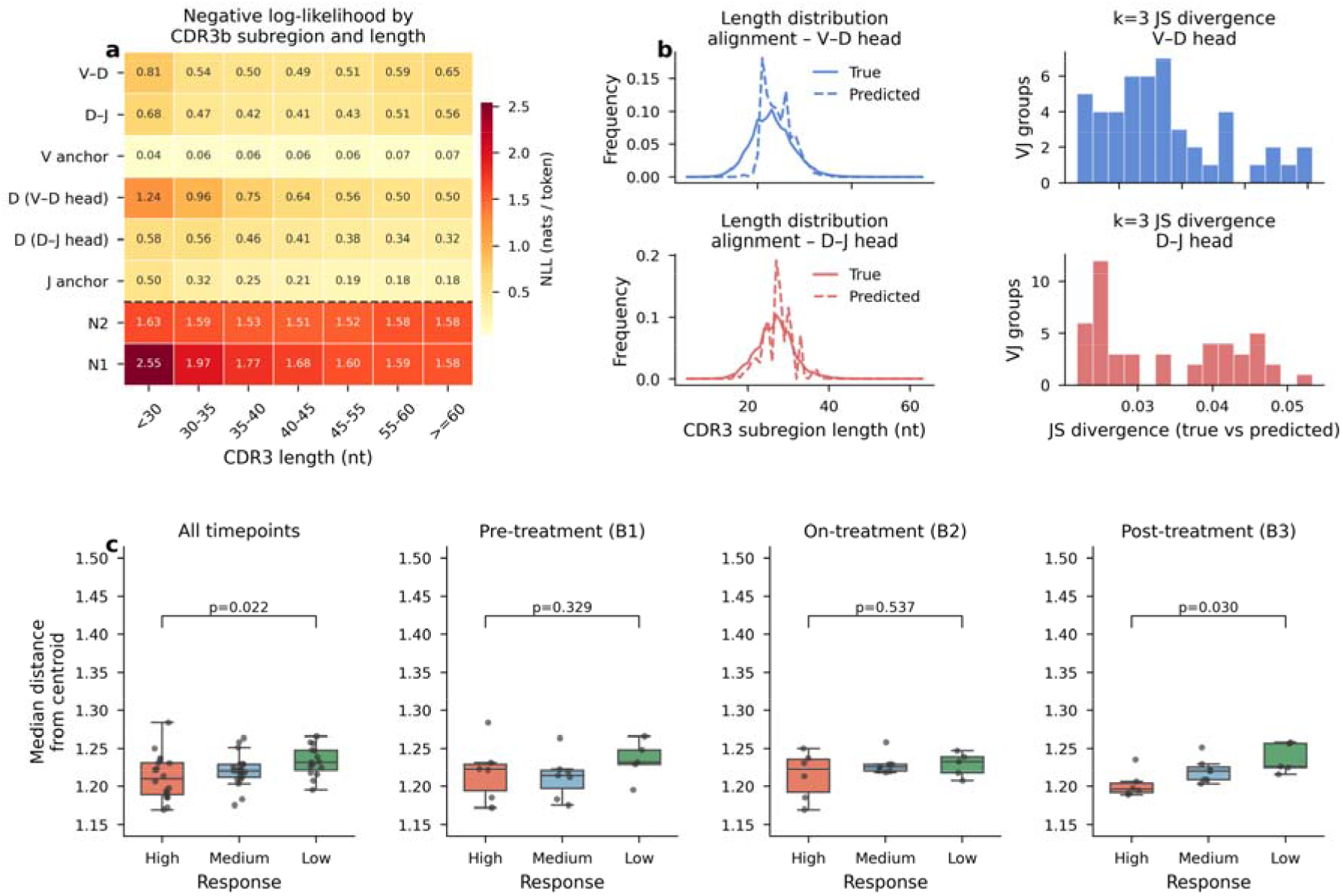
CRAFT captures response-associated repertoire organization in the OCSCC immunotherapy cohort. (a) Generative model performance: perplexity and NLL across CDR3-beta subregions (V-derived, N1, D-derived, N2, J-derived), showing lowest perplexity in germline-encoded regions and highest in N-insertion regions. (b) CRAFT reproduces empirical CDR3-beta length distributions and nucleotide k-mer composition across VJ strata, with low Jensen-Shannon divergence. (c) Distance from the centroid of a patient’s TCR repertoire in embedding space for the most expanded clones, stratified by response group and timepoint (B1, baseline; B2, on-treatment; B3, post-treatment).

### Dynamic reshaping of the TCR landscape during immunotherapy

We next evaluated whether CRAFT embeddings can characterize TCR repertoire structure and remodeling in cancer patients undergoing immunotherapy. Because CRAFT is trained exclusively on healthy-donor repertoires, deviation from its learned manifold provides a quantitative measure of immune perturbation. We analyzed a published cohort of 18 patients with oral cavity squamous cell carcinoma (OCSCC) treated with neoadjuvant checkpoint blockade.^21^ Patients received anti-PD-1 monotherapy (nivolumab) or combination anti-PD-1^22,23^ plus anti-CTLA-4^24^ (ipilimumab) therapy, with bulk PBMC TCR-beta sequencing at baseline (B1), on-treatment (B2), and post-treatment (B3) timepoints. Treatment arm and pathological response are partially confounded in this small cohort (Table S2), and these analyses should be interpreted as proof-of-concept demonstrations of the analytic framework.

We first examined local organization among dominant clonotypes by measuring the median distance of the top 100 clonotypes from each sample’s centroid. High responders showed more tightly clustered dominant clonotypes (p = 0.022, Figure 2c), and the separation was most prominent post-treatment (B3: p = 0.03), whereas differences at baseline and on-treatment were not significant (B1: p = 0.329; B2: p = 0.537), consistent with the possibility that treatment amplifies a pre-existing but initially undetectable organizational difference.

We next evaluated variance in the CRAFT embedding space; high responders showed lower variance (p = 0.022), most evident post-treatment (B3: p = 0.082; B2: p = 0.082; B1: p = 0.931; Figure S2). This lower variance suggests that the most expanded clones in responders share structural similarity in CDR3 sequence space. Biologically, a lower variance may indicate that the repertoire of responding patients is more focused in the embedding space, potentially reflecting coordinated clonal convergence toward shared antigenic targets rather than the dispersed clonal architecture seen in non-responders.

Longitudinally, high responders showed larger centroid shifts between baseline and post-treatment (Figure S2), consistent with more pronounced global repertoire remodeling during treatment. A large centroid shift indicates that the overall composition of the repertoire has moved substantially in the embedding space, consistent with therapy-induced clonal replacement or expansion of previously minor clonotypes.

### Compartment-specific longitudinal immune remodeling

We next analyzed matched single-cell TCR and RNA sequencing data from serial tumor biopsies of two patients with recurrent glioblastoma (GBM) treated with intratumoral oncolytic HSV-1 (CAN-3110)^25^. This analysis comprises only two patients and is intended to illustrate the resolution of CRAFT’s compartment-level framework in a richly annotated single-cell setting, not to support generalizable clinical conclusions.

Using a baseline-anchored framework, we computed clonotype distances from baseline centroids within three T cell compartments identified by scRNA-seq (using 20 PCs for single-cell data to avoid noise amplification; results were consistent across 5–40 PCs, Figure S4): CD8+ cytotoxic, CD8+ effector memory (EM), and stress-signature T cells (Figure 3a,b). Clonotypes exceeding the 90th-percentile baseline distance threshold were classified as “highly displaced,” and the fraction of displaced clonotypes was assessed at each timepoint using a binomial test (Figure 3c).

**Figure 3.**
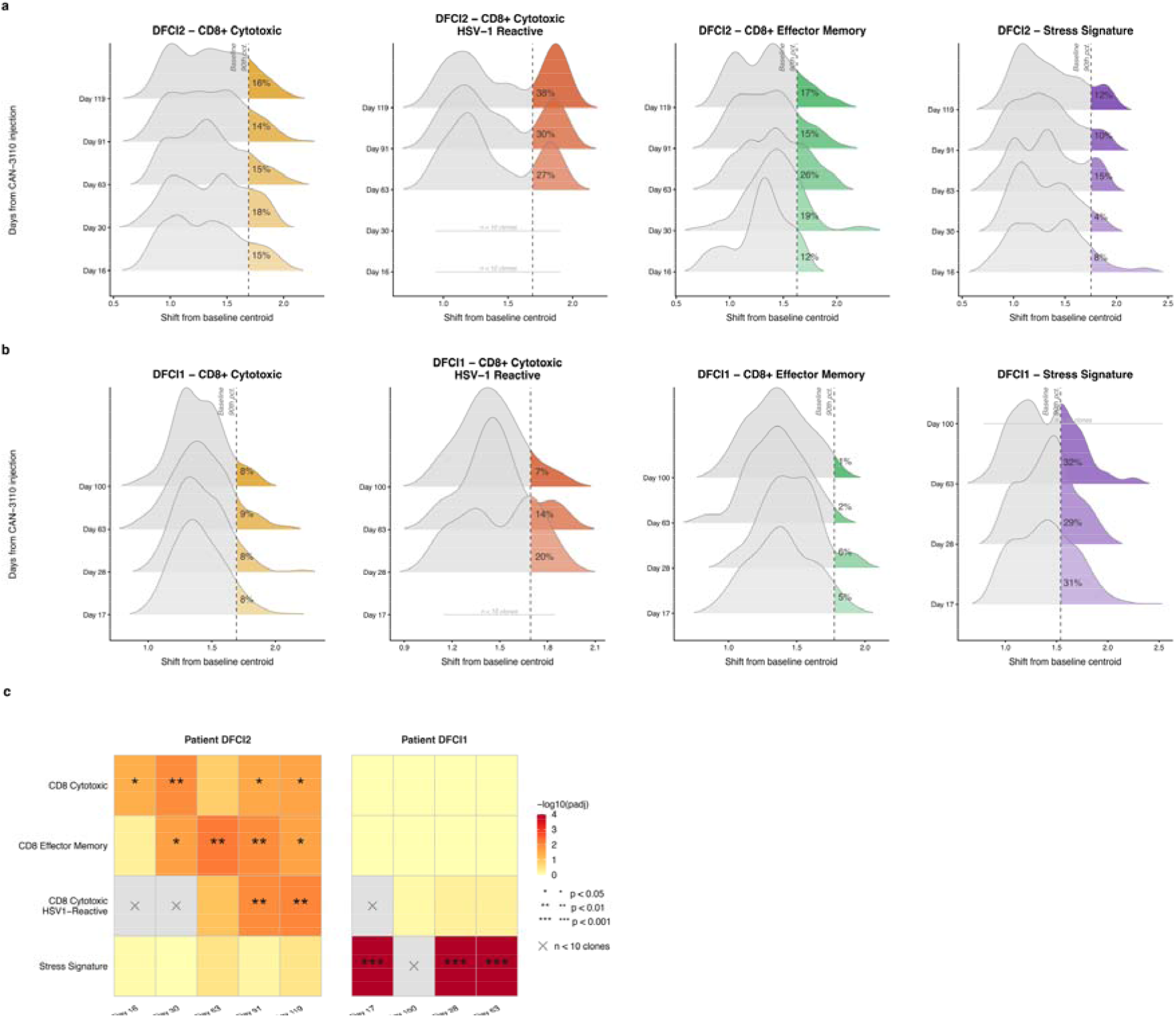
CRAFT embeddings capture compartment-specific longitudinal immune remodeling in recurrent BM. (a) Clonotype distance distributions from baseline centroid across timepoints for DFCI2 (clinically stable patient), shown for CD8+ cytotoxic T cells (all clonotypes and HSV-1-reactive), CD8+ effector memory T cells, and stress-signature T cells. Dashed line indicates the baseline 90th-percentile displacement threshold. (b) As in (a), for DFCI1 (progressing patient). (c) Binomial tail enrichment (fraction of displaced clones) by compartment and timepoint for both patients, with BH-adjusted p-values.)

The two patients exhibited divergent compartment-specific trajectories. In the clinically stable patient (DFCI2), CD8+ cytotoxic and effector memory T cell distance distributions broadened progressively from day 30 onward, with significant tail enrichment at multiple timepoints in both compartments (CD8+ cytotoxic: p_adj_ = 0.042, 0.010, 0.030, 0.025 at days 16, 30, 91, and 119; CD8+ EM: p_adj_ = 0.021–0.025 from day 30 onward; Figure 3c). Stratification by TCR reactivity revealed that this displacement was driven by HSV-1-reactive clonotypes, while EBV-reactive clonotypes remained near baseline (Supplementary Information), consistent with antigen-specific clonal expansion selectively reshaping the repertoire in response to the oncolytic virus. In contrast, the progressing patient (DFCI1) showed no systematic displacement in either CD8+ compartment; instead, the stress-signature compartment exhibited large clonal expansion at day 17 with sustained enrichment through day 63 (p_adj_ = 7.6 × 10^-31,^ 1.6 × 10^-4,^ 6.4 × 10^-6^; Figure 3c), a pattern consistent with progressive T cell dysfunction under persistent tumor pressure.

The contrasting patterns of antigen-driven cytotoxic displacement in the stable patient versus stress-associated expansion in the progressing patient were detectable only because the single-cell framework measures displacement within defined cellular compartments relative to an external healthy-donor reference. Such compartment-specific signals would be invisible in bulk analysis. Differential gene expression analysis comparing highly versus minimally displaced CD8+ cytotoxic clonotypes in the clinically stable patient (DFCI2) revealed enrichment for cytotoxic and possibly terminally differentiated markers (KLRD1, KLRG1, NKG7) in high-displacement clones and memory-associated markers (IL7R, ICOS) in low-displacement clones (Figure S3), providing orthogonal transcriptional evidence that geometric distances in the CRAFT space track biologically meaningful phenotypic transitions (Supplementary Information).

### Antigen-class structure in pathology-associated TCRs

Analysis of 5,279 pathology-associated TCRs from McPAS-TCR^26^ showed partial separation of antigen classes in the CRAFT embedding space (Figure 4a). Autoimmune- and pathogen-specific TCRs formed distinct clusters, with cancer-associated TCRs peripheral to the pathogen cluster. Using 500 balanced down-sampled replicates, silhouette scores were positive for all classes (allergy = 0.224, autoimmune = 0.066, cancer = 0.066, pathogens = 0.042; Figure 4b), and kNN class-consistency confirmed local neighborhood structure (pathogens = 0.886, autoimmune = 0.840, cancer = 0.731, allergy = 0.541; Figure 4d). Hierarchical clustering of class centroids grouped cancer and pathogen associated TCRs together, with allergy and autoimmune forming a separate branch (Figure 4c). The grouping of cancer with pathogen associated TCRs may reflect shared selective pressures, as both cancer neoantigens and viral antigens drive strong cytotoxic responses. These modest silhouette scores reflect the inherent limitations of CDR3-beta-only analysis; antigen recognition depends on the paired alpha-beta heterodimer, and CDR3-beta captures only a fraction of specificity information. A representative subset of UMAPs^27^ from different down-samples is shown in Figure S5.

**Figure 4.**
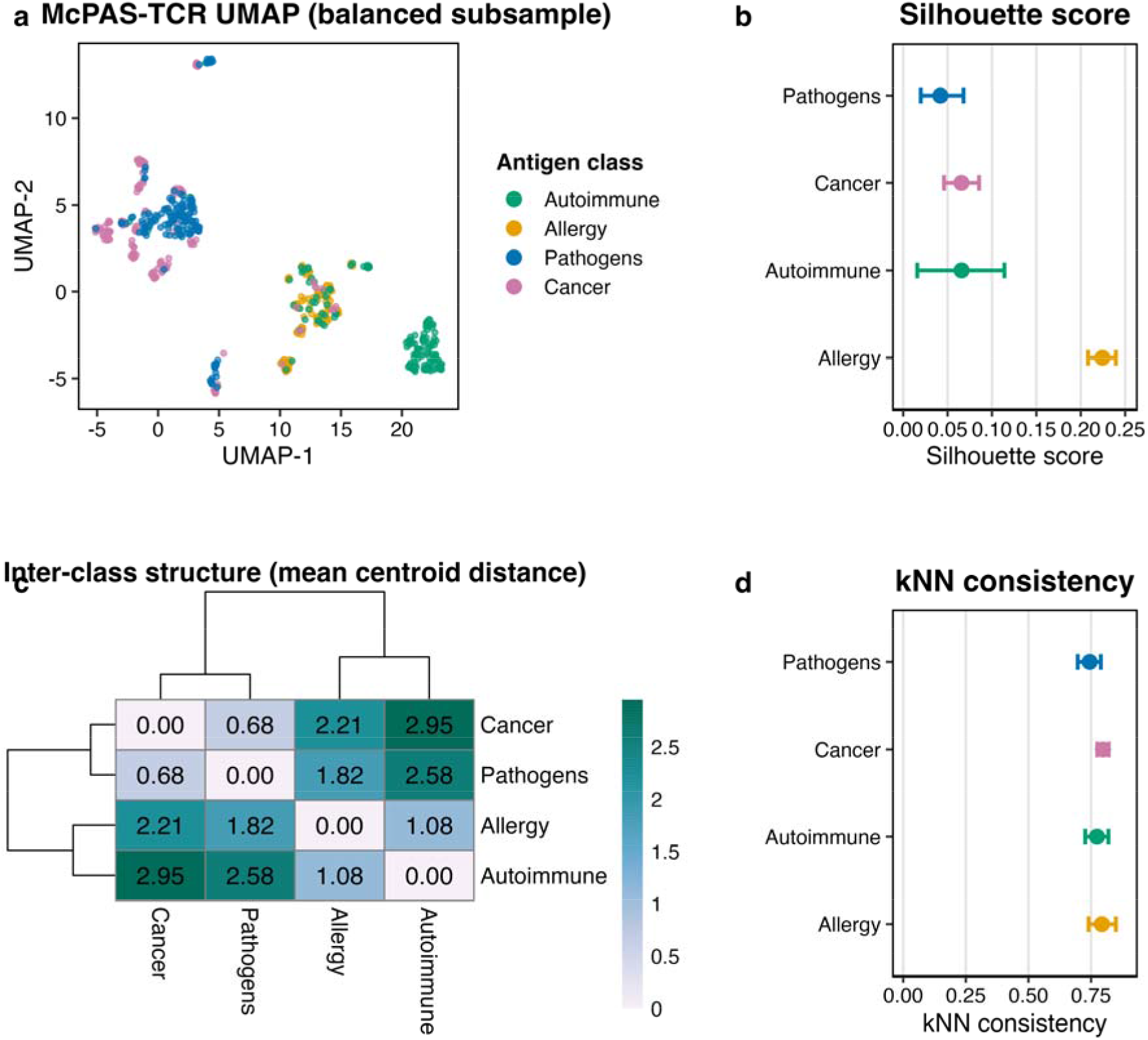
CRAFT embeddings capture antigen-class structure in pathology-associated TCRs from McPAS-TCR. (a) UMAP visualization of CRAFT embeddings for a balanced subsample (150 sequences per antigen class), colored by class (Pathogens, Autoimmune, Cancer, Allergy). (b) Per-class silhouette scores across 500 balanced replicates. (c) Pairwise centroid distances with hierarchical clustering, showing cancer-pathogen and allergy-autoimmune groupings. (d) kNN class-consistency scores (k = 10) across 500 balanced replicates.

### Comparison with other embedding methods

We benchmarked CRAFT against TCR-BERT^13^, GIANA^12^, SCEPTR^15^, and ESM^16^ across three evaluation settings (Figure 5). An important caveat applies to all comparisons: because CRAFT operates on beta-chain nucleotide sequences, all methods were evaluated using CDR3-beta input only. Three of the four comparators (ESM, TCR-BERT, and SCEPTR) natively support paired alpha-beta amino-acid input and restricting them to beta-chain sequences may understate their full capacity. The benchmarks, therefore, assess representation quality under a shared input modality, not each method’s best-case performance.

**Figure 5.**
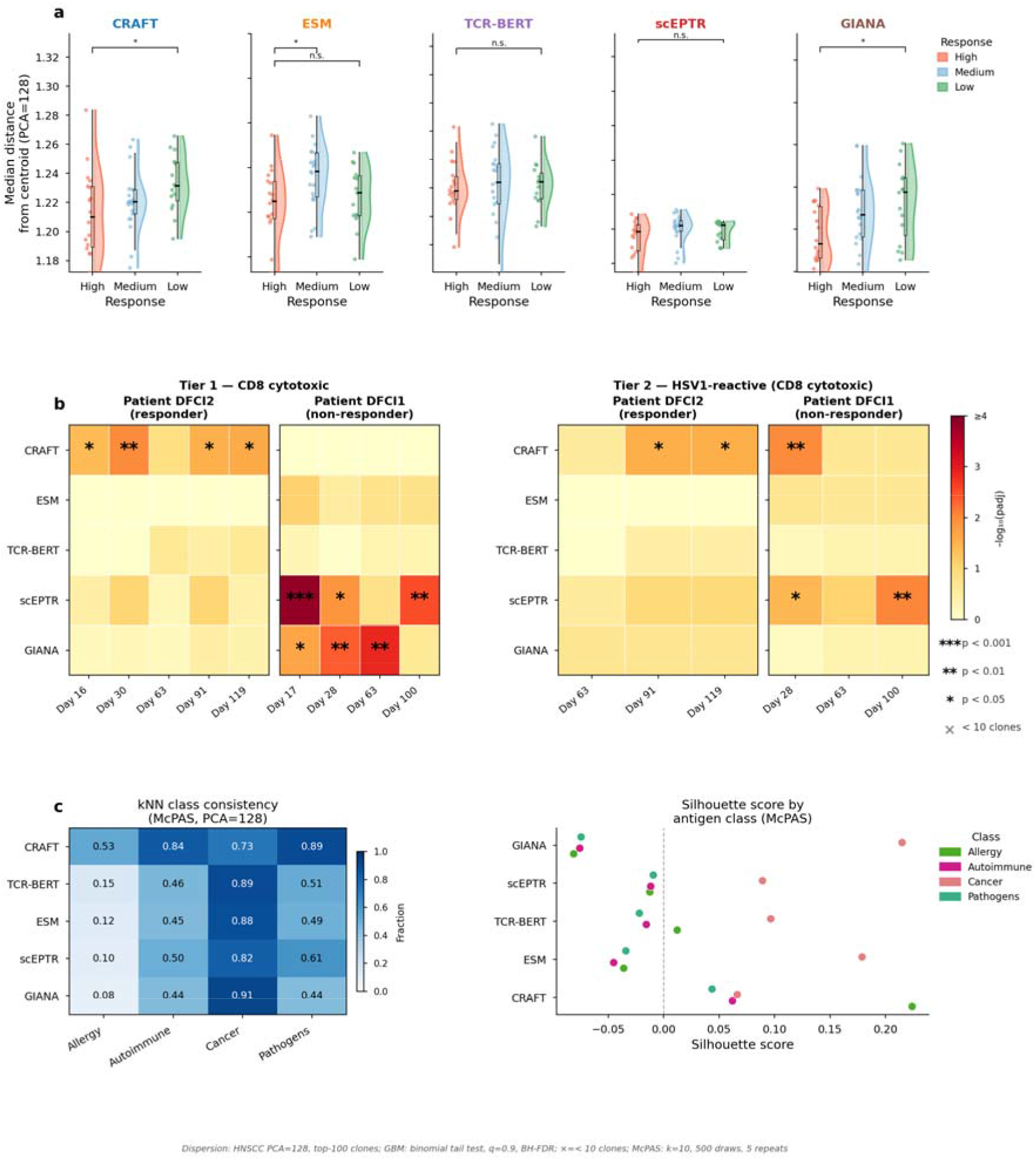
Comparison of CRAFT with other TCR embedding methods across three evaluation settings: (a) global immune remodeling (OCSCC cohort median distance from centroid by response group), (b)compartment-specific remodeling (GBM cohort, displacement-based tail enrichment in CD8+ cytotoxic compartment across methods), and antigen-class structure (McPAS-TCR, silhouette scores and kNN consistency across methods).

#### Global immune remodeling

In the OCSCC cohort (n=18), only CRAFT and GIANA significantly distinguished response groups by median distance from centroid when pooling across timepoints (CRAFT: p = 0.022; GIANA: p = 0.013; Figure 5a). These comparisons are drawn from the same 18-patient cohort and should be interpreted with caution given the limited statistical power. Notably, the two methods showed complementary temporal profiles: CRAFT achieved its clearest separation post-treatment (B3: p = 0.03), whereas GIANA showed its strongest signal at baseline (B1: p = 0.082). This temporal distinction is informative: CRAFT’s post-treatment signal is consistent with detection of therapy-induced repertoire remodeling, the clonal replacement and expansion that accumulate over the course of immunotherapy, whereas GIANA, a sequence-similarity clustering method, may be more sensitive to pre-existing clonal structure that is already apparent early in treatment. A similar pattern emerged for repertoire variance across all clonotypes: high responders showed lower variance in embedding space, and again only CRAFT and GIANA significantly distinguished response groups when pooling across timepoints (CRAFT: p = 0.022; GIANA: p = 0.0016). The temporal profiles again diverged: CRAFT showed a trend toward separation at both on- and post-treatment timepoints (B2 and B3: p = 0.082 each; B1: p = 0.931), whereas GIANA’s strongest separation occurred on-treatment (B2: p = 0.017), further supporting the interpretation that CRAFT preferentially captures therapy-driven repertoire changes.

#### Compartment-specific remodeling

We next benchmarked compartment specific remodeling in the two GBM patients. In the clinically stable patient (DFCI2), CRAFT was the only method to detect statistically significant enrichment of displaced CD8+ cytotoxic clones at multiple timepoints (Days 16, 30, 91, and 119; p_adj_ = 0.042, 0.010, 0.030, and 0.025 respectively, Figure 5b), and all other methods, including ESM, TCR-BERT, SCEPTR, and GIANA, failed to detect enrichment at any timepoint in this patient. Notably, GIANA and SCEPTR, each detected enrichment at three timepoints in the progressing patient (DFCI1) but produced no significant signal in DFCI2, suggesting these methods may capture a different axis of clonal variation unrelated to antigen-driven displacement. Restricting to HSV1-reactive clones, CRAFT again uniquely identified significant enrichment at Days 91 and 119 (p_adj_ = 0.029), while no other method reached significance for this subpopulation. A Fisher’s exact test comparing HSV1 versus EBV tail fractions at each timepoint was underpowered due to the small number of EBV-reactive clones (n <= 10 per timepoint) and yielded no method-timepoint combination surviving BH correction, though CRAFT consistently showed the largest directional effect (HSV1 tail fraction 30-38% versus EBV 6-13% at Days 91-119). Taken together, these results indicate that CRAFT’s embeddings confer substantially greater sensitivity for detecting antigen-driven clonal displacement than sequence-similarity or language-model baselines, and that the HSV1-reactive enrichment signal is specific to CRAFT’s representation geometry.

#### Antigen-class structure

When restricted to CDR3-beta input alone, CRAFT achieved positive silhouette scores across all four antigen categories, whereas other methods showed higher cancer-class scores but negative scores for the remaining categories. CRAFT also showed higher kNN consistency for non-cancer classes (e.g., pathogens: 0.88 vs. 0.43 for GIANA, Figure 5c). As noted above, all methods were constrained to beta-chain input for these comparisons.

## Discussion

CRAFT learns a reference grammar of productive TCR-beta generation from healthy individuals and uses it as a coordinate system for characterizing cancer-associated repertoires as structured departures from baseline. This framing repurposes the embedding as a quantitative substrate for measuring immune perturbation and remodeling at the repertoire level, a different goal from the epitope classification that most TCR embeddings are optimized for. The most informative signals in our analyses were distributional: geometric changes across many clonotypes in latent space, including dispersion, centroid organization, global movement, and local density. This is conceptually appealing because immunotherapy response involves systems-level reweighting of clonal frequencies and phenotypic transitions across the repertoire. Lower embedding variance in responders indicates a more focused, coordinated repertoire; larger centroid shifts reflect wholesale repertoire remodeling; and tail enrichment of displaced clonotypes pinpoints compartments and timepoints where antigen-driven selection is strongest. Because the embedding is defined relative to a reference grammar rather than fitted per dataset, these patient-level geometric shifts are comparable across cohorts and serve as interpretable readouts.

The single-cell glioblastoma analysis shows how a baseline-anchored reference resolves signals that bulk analysis would miss. Measuring displacement within defined T-cell compartments relative to an external healthy reference separated two biologically distinct trajectories: antigen-driven displacement of CD8+ cytotoxic cells in the clinically stable patient (DFCI2) and dysfunction-associated expansion of stress-signature cells in the progressing patient (DFCI1). That geometric distance in CRAFT space aligned with effector-differentiation markers (Figure S3) gives orthogonal transcriptional support that these distances track meaningful phenotypic transitions.

In head-to-head comparisons, CRAFT and GIANA most consistently recovered response-associated remodeling in the bulk OCSCC setting, possibly because both encode repertoire-structural information beyond purely protein-language features. In the single-cell GBM setting, CRAFT’s embeddings showed uniquely sustained sensitivity to antigen-driven displacement, whereas GIANA and SCEPTR detected signal preferentially in the progressing patient. Embeddings trained to represent the rules of repertoire formation may therefore be particularly suited to detecting antigen-specific change, while sequence-similarity methods may capture a broader, less specific axis of clonal variation. The natural next step is paired alpha-beta modeling, which should improve antigen-level resolution for specificities where alpha-chain contributions dominate.

## Limitations of the study

This work is a proof of concept with several limitations. The clinical analyses rest on small patient numbers: a checkpoint-blockade cohort of 18 OCSCC patients and a two-patient single-cell study of oncolytic immunotherapy in recurrent glioblastoma. Both were sampled longitudinally, the OCSCC cohort by bulk PBMC TCR-beta sequencing at three timepoints (baseline, on-treatment, post-treatment) and the two GBM patients by serial single-cell sequencing across five and six biopsies with T-cell compartments resolved separately, so each study contributes many observations across time and cellular compartments. The number of individuals nonetheless remains limited, treatment arm and pathological response are partly confounded in the OCSCC cohort (Table S2), and the response-associated geometric signals we report require validation in larger, independent cohorts with uniform regimens before clinical interpretation. CRAFT models the TCR-beta chain at the CDR3 and germline level and does not incorporate the alpha chain or paired-chain information, which limits resolution of antigen specificity; consistent with this, CDR3-beta alone provided only partial signal in several antigen-labeled classes. The healthy reference derives from a single large bulk-sequencing dataset; although CMV serostatus does not dominate the embedding space (Figure S1), broader demographic and technical diversity in the reference, together with conditioning on covariates such as age, infection history, and HLA, would strengthen generalization and will require reference cohorts with richer clinical annotation than are currently public. Finally, the geometric metrics quantify repertoire structure and its movement but do not by themselves establish the antigenic or functional drivers of the shifts they detect.

## STAR Methods

### KEY RESOURCES TABLE

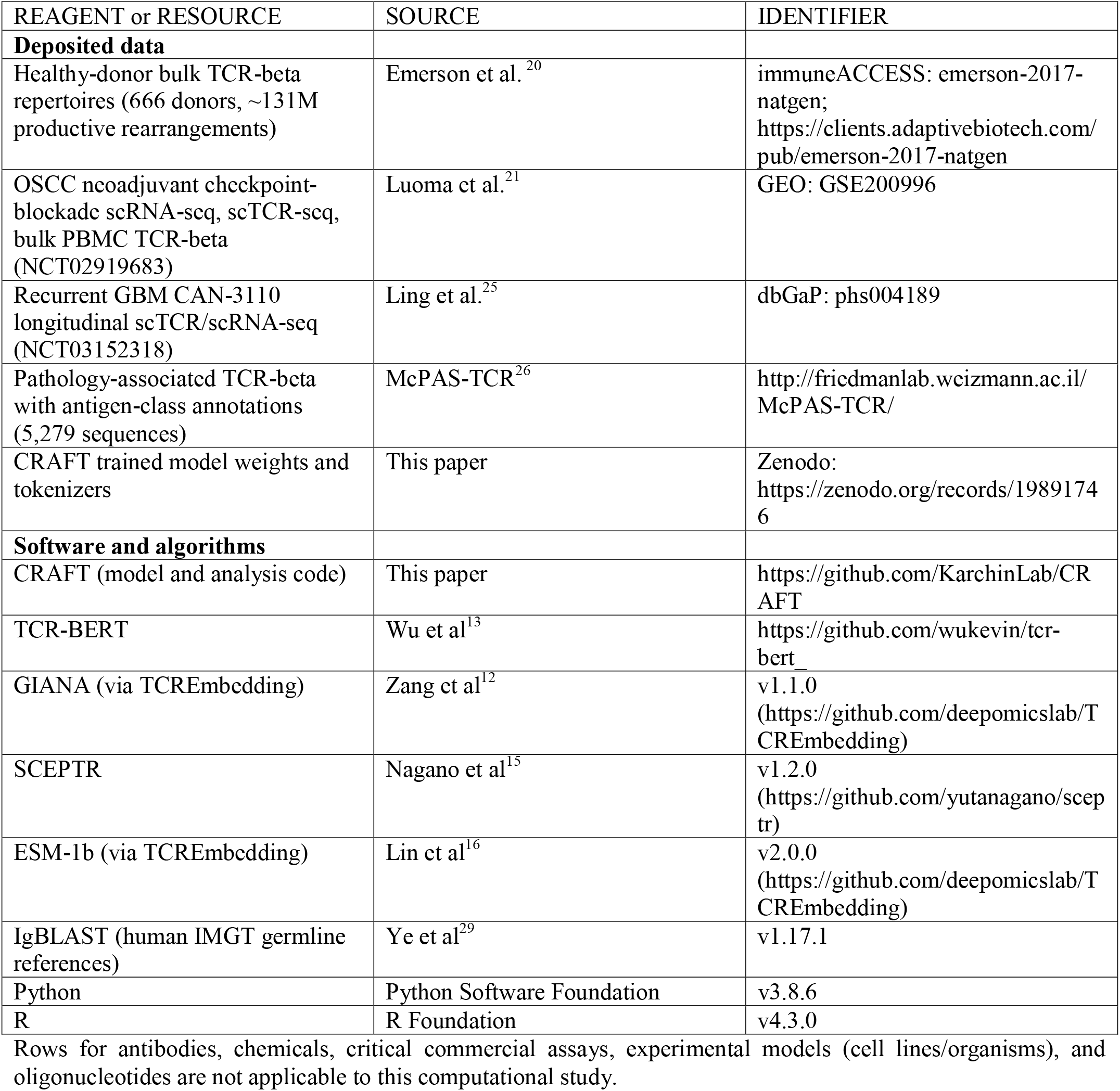

Rows for antibodies, chemicals, critical commercial assays, experimental models (cell lines/organisms), and oligonucleotides are not applicable to this computational study.

### RESOURCE AVAILABILITY

#### Lead contact

Further information and requests for resources should be directed to and will be fulfilled by the lead contact, Rachel Karchin.

#### Materials availability

This study did not generate new unique reagents.

#### Data and code availability

- This paper analyzes existing, publicly available data. The bulk TCR-beta repertoires from 666 healthy donors are available through the Adaptive Biotechnologies immuneACCESS repository (Emerson et al.^20^). The oral cavity squamous cell carcinoma neoadjuvant immunotherapy data (Luoma et al.^21^; NCT02919683) are available in the Gene Expression Omnibus under accession GSE200996. The recurrent GBM CAN-3110 single-cell data (Ling et al.^25^; NCT03152318) are available in the Database of Genotypes and Phenotypes under accession phs004189. Pathology-associated TCR annotations are available in the McPAS-TCR database. These accession numbers are also listed in the key resources table.
- All original code has been deposited at GitHub (https://github.com/KarchinLab/CRAFT) and is publicly available as of the date of publication. Trained model weights and tokenizers are deposited at Zenodo (https://zenodo.org/records/19891746). DOIs are listed in the key resources table.
- Any additional information required to reanalyze the data reported in this paper is available from the lead contact upon request.

### EXPERIMENTAL MODEL AND STUDY PARTICIPANT DETAILS

This study used only previously published, de-identified human data; no new human participants were enrolled and no new samples were collected. Detailed descriptions of all datasets are provided in Table S1, four datasets were used: (1)The Emerson et al.^20^ cohort of bulk peripheral-blood TCR-beta repertoires from 666 healthy donors (approximately 131 million productive rearrangements) was used for model training, with an 80/10/10 train/validation/test split at the sequence level. The validation cohort of 120 donors from the same study was used to test for batch effects in CRAFT embeddings. (2) The Luoma et al.^21^ oral cavity squamous cell carcinoma (OCSCC) cohort (NCT02919683) provided bulk PBMC TCR-beta sequencing from 18 patients receiving neoadjuvant checkpoint blockade at three timepoints. (3) The Ling et al.^25^ GBM cohort (NCT03152318) provided longitudinal single-cell TCR and RNA sequencing from serial tumor biopsies of two patients treated with intertumoral oncolytic HSV-1 (CAN-3110). (4) The McPAS-TCR^26^ database provided 5,279 human pathology-associated TCR-beta sequences with antigen-class annotations.

### METHOD DETAILS

#### TCR sequencing preprocessing

Bulk data (Adaptive immunoSEQ platform): Productive, in-frame rearrangements with resolved V(D)J assignments and valid V-gene annotations were retained. V, D, and J gene calls were standardized to IMGT format and decomposed into family, subfamily, and allele components. CDR3 nucleotide sequences were extracted and segmented into five subregions (V-derived, N1, D-derived, N2, J-derived) using Adaptive-provided positional indices. Composite recombination contexts (V-D, D-J, V-N1-D, D-N2-J) were constructed for model input.

Single-cell data: Single-cell TCR sequencing datasets lacking explicit positional indices were realigned using IgBLAST with human IMGT germline references to reconstruct recombination boundaries in a germline-aware manner. IgBLAST AIRR outputs were converted into the unified CRAFT input schema by extracting aligned V-, D-, and J-derived subsegments and NP1/NP2 junctional insertions. Sequences lacking valid V- or J-derived subsegments after reconstruction were excluded.

#### Model architecture and training

CRAFT is a conditional sequence-to-sequence transformer based on the BART architecture^17^. The encoder takes V, D, and J genes as inputs using a hierarchical IMGT-based gene tokenizer; the decoder uses character-level nucleotide tokenization (A, C, G, T) with two parallel output heads for the V-D and D-J junctions. Both heads share decoder hidden states and generate subregion-specific nucleotide predictions.

The model was trained using token-level cross-entropy loss applied to autoregressive CDR3 reconstruction. The total loss is a weighted sum of V-D and D-J head losses:

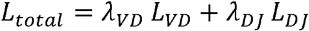

where each head loss is the negative log-likelihood averaged over sequence positions:

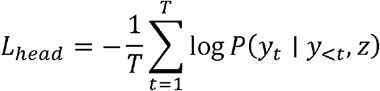

with *T* the sequence length, *y*_*t*_ the nucleotide at position *t, y*_<*t*_ the preceding nucleotides, and *z* the encoder representation of the germline context. Perplexity, the exponentiated average negative log-likelihood, was used as the primary evaluation metric:

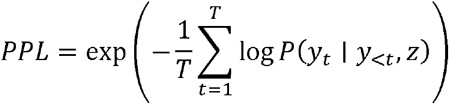

Optimization used AdamW with decoupled weight decay, a cosine learning-rate schedule with linear warmup, gradient clipping (max norm = 1.0), and dropout for regularization. Curriculum training proceeded through multiple phases of progressively increasing sequence complexity: early phases restricted training to shorter CDR3 sequences with common V-J pairings to establish stable gradient dynamics, while later phases introduced the full diversity of CDR3 lengths and gene combinations. Phase transitions were triggered after convergence on validation loss, defined as fewer than 0.1% improvement over 5 consecutive evaluation intervals. All hyperparameters for each curriculum phase are reported in Table S1.

#### Embedding extraction

Sequence embeddings were obtained using teacher-forced inference^18,19^, in which true V, D, J gene tokens and CDR3 subregion tokens were provided to the model. Hidden states from the V-D and D-J decoder branches were mean-pooled across tokens and concatenated to form a unified per-TCR representation for all downstream analyses.

Batch effects in the training data. To confirm that CRAFT embeddings reflect TCR sequence features rather than donor-level covariates, we computed mean-pooled embeddings per sample from the Emerson et al. validation cohort (n=120) and examined the top principal components for structure attributable to CMV status, age, or sex. No dominant covariate signal was observed in the top PCs (Figure S1).

#### Principal component analysis

PCA was applied to pooled CRAFT embeddings within each cohort for dimensionality reduction. The number of retained PCs was chosen based on sample size relative to embedding dimensionality: for bulk analyses with large per-sample clonotype counts (OCSCC) and sequence-level antigen analyses (McPAS-TCR), 128 PCs were retained; for single-cell analyses with smaller per-compartment counts (GBM), 20 PCs were retained to avoid noise amplification in low-count settings.

To verify that downstream conclusions are not sensitive to this choice, we repeated all primary analyses across a range of PC dimensions (5, 10, 20, 30, and 40 PCs). Effect directions were consistent across all settings: response-group separation in the OCSCC cohort, compartment-specific tail enrichment in the GBM cohort, and antigen-class clustering in McPAS-TCR all showed the same qualitative patterns regardless of the number of PCs retained. Statistical significance was preserved at all tested dimensions for the primary findings. These results confirm that the reported signals reflect stable structure in CRAFT embeddings rather than artifacts of a particular dimensionality-reduction setting (Figure S4).

#### Bulk TCR repertoire metrics

All metrics below operate on PCA-projected CRAFT embeddings. Let *x*_*i*_ ∈ℝ ^*d*^ denote the embedding of clonotype *i* in sample *s* containing *n*_*s*_ clonotypes. Embedding variance. The sample mean is:

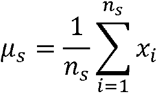

The sample covariance matrix is:

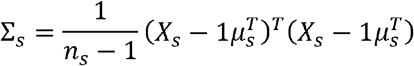

where *X*_*s*_ is the *n*_*s*_ ⨯ *d* embedding matrix and 1 is a column vector of ones. Embedding variance is:

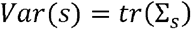

This scalar summarizes the total dispersion of the repertoire in latent space.

Median distance from centroid. For the top *N* clonotypes by frequency (*N =* 100):

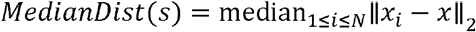

where *x* is the centroid of the top *N* clonotypes. This measures how tightly the dominant clonotypes cluster in latent space.

Centroid shift. For patient *p* between timepoints *t*_1_ and *t*_2_:

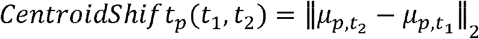

where *μ*_*p,t*_ is the L2-normalized sample centroid at timepoint *t*. is the L2-normalized sample centroid at timepoint between timepoints.

#### Single-cell TCR metrics

For each patient, cell type, and timepoint, single-cell embeddings were aggregated to the clonotype level by computing the mean embedding across all cells of the same clonotype.

Baseline centroid (weighted). At the earliest timepoint *t*_0_ with at least 20 clonotypes, a weighted baseline centroid is defined as:

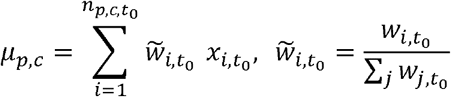

where *w*_*i*_ is the number of cells contributing to clonotype *i* at baseline.

Distance to baseline. For each clonotype *j* at timepoint *t*:

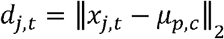

Baseline tail threshold. The 90th percentile of baseline distances defines the displacement threshold:

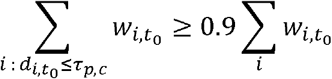

Clonotypes with *d*_*j*_,_*t*_ > *τ*_*p,c*_ at subsequent timepoints are classified as “highly displaced.”

Weighted tail mass. At each timepoint *t*, the weighted tail mass is the fraction of total cellular mass contributed by displaced clonotypes:

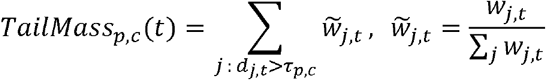

where *W*_*jt*_ is the number of cells for clonotype *j* at timepoint *t*. Statistical significance of tail enrichment was assessed using a binomial test against the expected baseline tail fraction (0.10).

For reactivity-stratified analyses, baseline centroids and thresholds were computed using all clonotypes irrespective of reactivity, and weighted tail mass was computed separately within each reactivity class. For differential expression, Wilcoxon rank-sum tests were applied to log-normalized counts comparing cells from clones in the top versus bottom 25th percentile of CRAFT displacement from baseline within each timepoint (Figure S3).

#### Antigen specificity metrics

To account for class imbalance, all evaluations used 500 balanced downsampled replicates (*N*_*max*_ =150 per class).

Silhouette score. For each sequence *i* with class label *y*_*t*_:

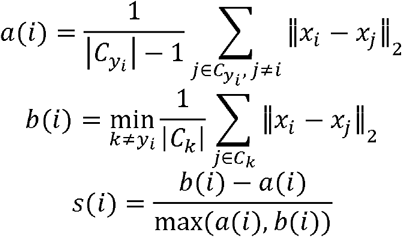

where *a*(*i*) is the mean intra-class distance, *b*(*i*) is the mean distance to the nearest other class, and *C*_*k*_ denotes the set of sequences in class *k*. Per-class silhouette scores are averaged over all sequences in each class.

kNN class-consistency. For each sequence *i* with *k =* 10 nearest neighbors:

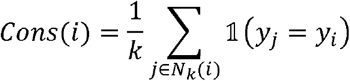

where 1 is the indicator function. This measures the fraction of nearest neighbors sharing the same class label, quantifying local neighborhood purity. Pairwise Euclidean distances between antigen-class centroids were computed and used for hierarchical clustering. All metrics were averaged across the 500 replicates.

#### Benchmark embedding methods

TCR-BERT^13^ was applied to CDR3-beta amino-acid sequences using the pretrained model from the original publication. SCEPTR^15^ used CDR3-beta amino-acid sequences with V and J gene annotations. GIANA^12^ was applied to CDR3-beta amino-acid sequences using the TCREmbedding package^28^. ESM embeddings were generated using the ESM-1b model^16^ (esm1b_t33_650M_UR50S) applied to CDR3-beta amino-acid sequences. For all methods, gene nomenclature was standardized to IMGT format, and sequences were filtered to remove noncanonical characters and sequences below minimum length thresholds. Embeddings were projected using PCA and evaluated with the same downstream metrics applied to CRAFT.

### QUANTIFICATION AND STATISTICAL ANALYSIS

All statistical analyses were performed in Python (v3.8.6) and/or R (v4.3.0). Tests were two-sided at alpha = 0.05. Nonparametric tests were used throughout given modest sample sizes and non-normal distributions. Group comparisons on patient-level CRAFT-derived metrics used the Mann-Whitney U test. For longitudinal analyses, metrics were computed per patient prior to group comparison, ensuring the patient was the unit of inference. Where multiple hypotheses were tested within the same analysis family, p-values were adjusted using Benjamini-Hochberg FDR correction (q < 0.05), and both nominal and adjusted p-values are provided. Summary statistics are reported as median and interquartile range. Exact p-values are reported when available (values below the reporting limit shown as p < 0.001). Robustness analyses were assessed by repeating full statistical comparisons across parameter settings and verifying consistency in effect direction and significance. Statistical details for each analysis, including the test used, n, and the definition of the unit of inference, are also given in the relevant figure legends.

## Acknowledgments

***Break Through Cancer*, Linghua Wang:** L.W. reports grant funding from Break Through Cancer. **Adrienne Boire:** Supported by NCI Cancer Center Support Grant 5P30CA008748-57. **Vinay K. Puduvalli:** Break Through Cancer. **Charles A. Whittaker:** Partially supported by NCI Cancer Center Support (Core) Grant P30-CA14051 to the Koch Institute. **Viviane Tabar:** Supported by NCI Cancer Center Support Grant P30 CA008748. **Michael J. Cima:** Break Through Cancer. **David A. Reardon:** Break Through Cancer GBM TeamLab, **Rachel Karchin:** R.K. reports grant funding from Break Through Cancer.

## Author contributions

**Conceptualization:** Archana Balan, Rachel Karchin, Betty Y.S. Kim, Pratibha Sharma. **Data Curation:** Archana Balan, Kevin E. Meza Landeros, Calixto-Hope G. Lucas, Gerard Baquer, Syed Shujaat Ali Zaidi. **Funding Acquisition:** Rachel Karchin, Michael J. Cima. **Investigation:** Archana Balan, Rachel Karchin, Yuval Elhanati, E. Antonio Chiocca, Betty Y.S. Kim, Adrienne Boire, Vinay K. Puduvalli, Alexei Stortchevoi, David A. Reardon. **Methodology:** Archana Balan, Rachel Karchin, Yuval Elhanati, E. Antonio Chiocca, Kenny Kwok Hei Yu, Alexei Stortchevoi. **Project Administration:** Rachel Karchin, Kenny Kwok Hei Yu, Vinay K. Puduvalli, Jennifer Gantchev, David A. Reardon. **Resources:** Rachel Karchin, E. Antonio Chiocca, Calixto-Hope G. Lucas, Syed Shujaat Ali Zaidi, Charles A. Whittaker, Alexei Stortchevoi, Pratibha Sharma, Viviane Tabar, David A. Reardon. **Software:** Archana Balan **Supervision:** Rachel Karchin, Calixto-Hope G. Lucas, Vinay K. Puduvalli. **Writing – Original Draft:** Archana Balan, Rachel Karchin, Valsamo Anagnostou. **Writing – Review and Editing:** Mesut Unal, Linghua Wang, E. Antonio Chiocca, Betty Y.S. Kim, Calixto-Hope G. Lucas, Emerenziana Runco, Adrienne Boire, Forest M. White, Jiaying Lai, David A. Reardon, Marcus De Almeida Mendes, Dimitrios N. Sidiropoulos.

## Data Science Teamlab Members

Alex K. Shalek, Andrew McPherson, Benjamin Haibe-Kains, Caroline Chung, Cheng-Zhong Zhang, Christina Curtis, Christine A. Pratilas, Elana Fertig, Ethan Cerami, Rachel Karchin, Rameen Beroukhim, Wesley Tansey, Afrooz Jahedi, Bethany C. Taylor, Charlie Whittaker, Christopher Tosh, Emma Dyer, Greg Raskind, Jake June-Koo Lee, Jovana Pavisic, Kimal Rajapakshe, Linghua Wang, Long Yuan, Luciane T. Kagohara, Manuel Schuerch, Shahab Sarmashghi, Siri Palreddy, Sophie Webster, Stuart Levine

## Accelerating GBM Therapies TeamLab Members

David A. Reardon, E. Antonio (Nino) Chiocca, Keith Ligon, Franziska Michor, Rameen Beroukhim, Adrienne Boire, Andrew McPherson, Bethany C. Taylor, Kenny Yu, Viviane Tabar, Betty Kim, Padmanee Sharma, Shiao-Pei Weathers, Vinay K. Puduvalli, Jian Hu, Kadir Akdemir, Sangeeta Goswami, Sreyashi Basu, Wen Jiang, Mesut Unal, Chetan Bettegowda, Calixto-Hope Lucas, Christopher Douville, Connor J. Liu, Long Yuan, Nathalie Y.R. Agar, C. Zoe Linke, Forest White, Michael J. Cima, Stuart Levine, Marcus Borenäs, Joshua D. Bernstock, Emma Dyer, Jennifer Gantchev, Jaimie Lee, Manuel Schuerch, Shahab Sarmashghi, Siri Palreddy, Chris Jannotta

## Declaration of generative AI and AI-assisted technologies in the manuscript preparation process

During the preparation of this work, the authors used Claude (Anthropic) to assist with copyediting and manuscript formatting, code generation, plus code repository organization and management. The authors reviewed all results and edited the content as needed and take full responsibility for the content of the publication.

## Supplemental information

**Figure S1.**
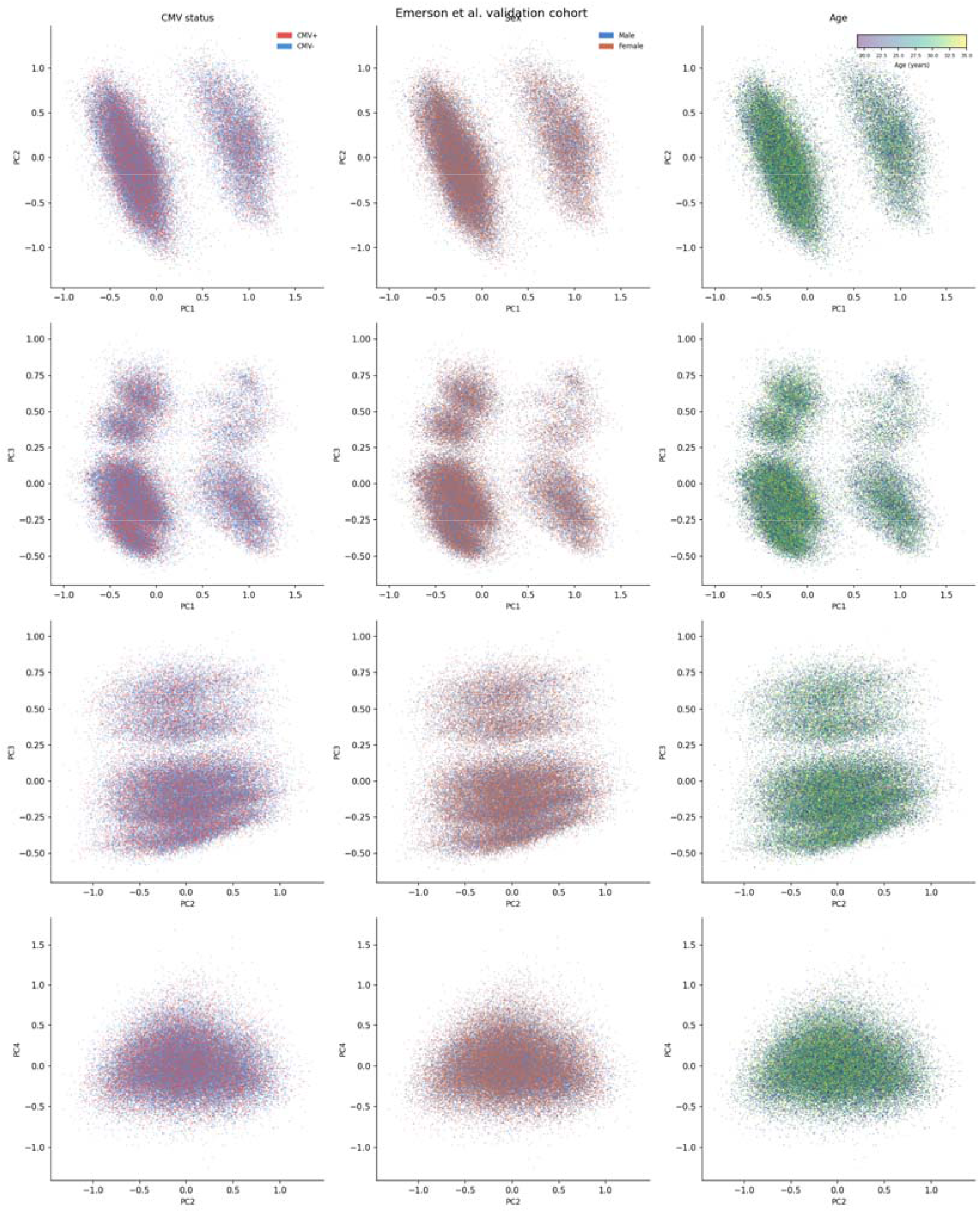
CMV serostatus does not dominate the CRAFT embedding space. PCA of per-donor embedding centroids for the 120 healthy donors in the Emerson et al. validation cohort, colored by CMV serostatus (positive vs. negative). Because CMV is among the strongest known drivers of TCR repertoire structure, we assessed whether CMV status organizes the learned representation. The inclusion of both CMV+ and CMV-donors in training is a deliberate design choice: the baseline reference should encompass the full range of healthy repertoire variation, including the substantial reshaping driven by chronic viral exposures, so that deviations measured in cancer cohorts reflect perturbations beyond this normal range.

**Figure S2.**
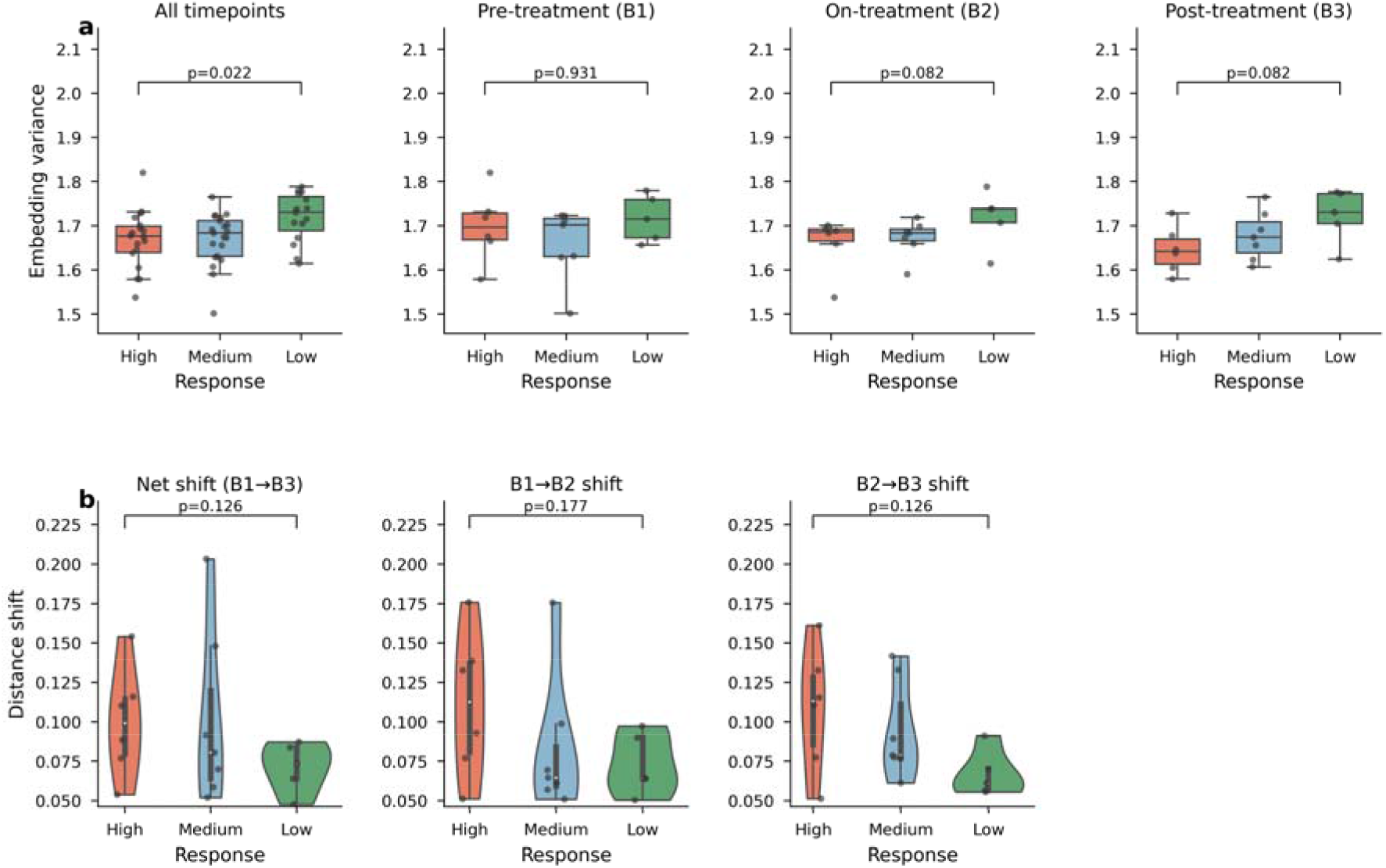
OCSCC cohort (a) embedding variance and (b) trends of longitudinal shifts

**Figure S3.**
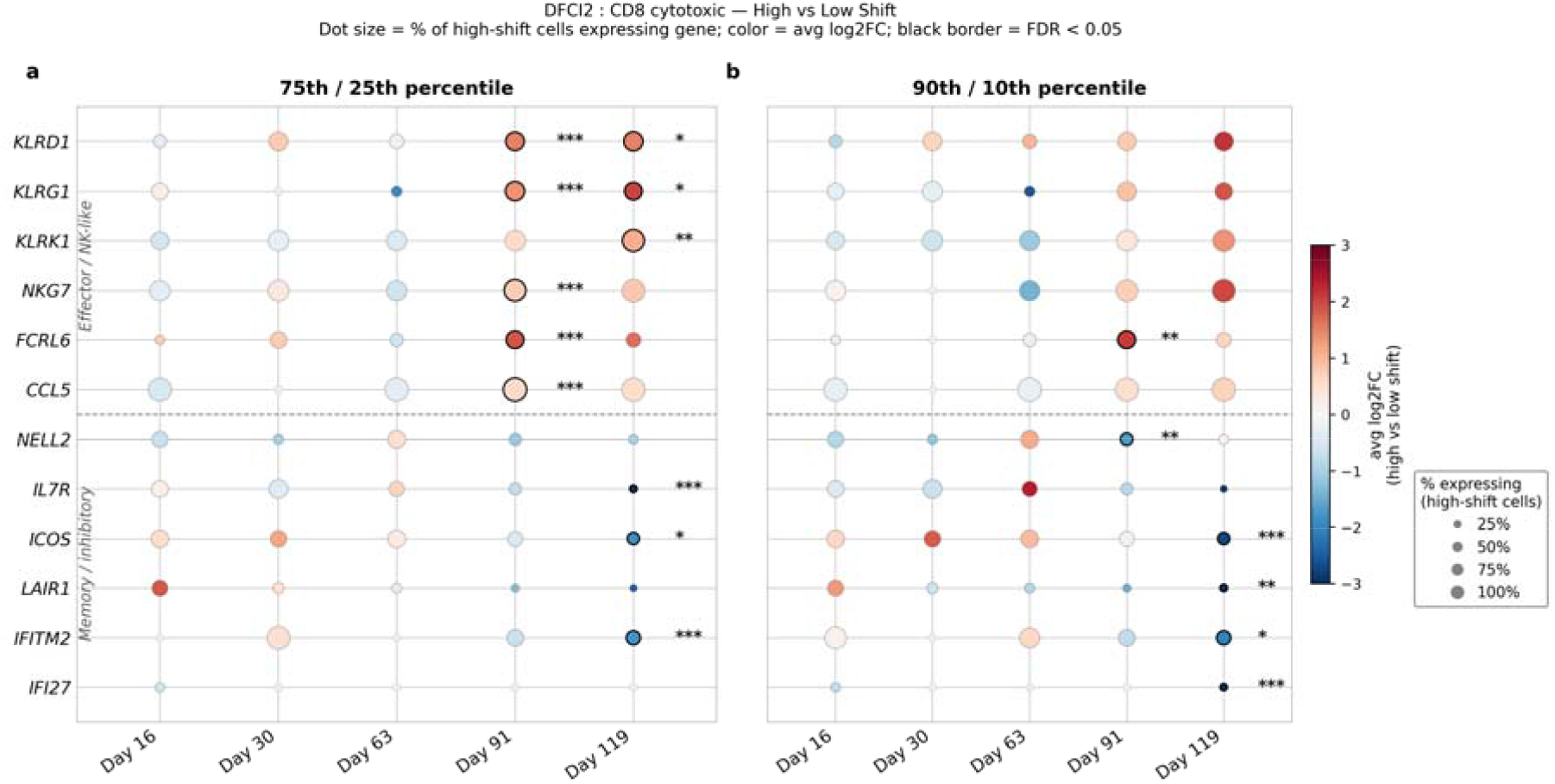
Differential gene expression between highly displaced and minimally displaced clonotypes in the GBM dataset. For compartments that showed statistically significant tail enrichment (DFCI2 CD8+ cytotoxic T cells, DFCI2 CD8+ effector memory T cells, and DFCI1 stress-signature T cells), single-cell differential expression analysis compared cells from clones in the top versus bottom 25th percentile of CRAFT displacement from the baseline centroid at each timepoint using a Wilcoxon rank-sum test on log-normalized counts. In DFCI2 CD8+ cytotoxic T cells, high-shift clones at day 91 were enriched for markers of terminal effector differentiation and NK-like cytotoxic function, including KLRD1 (CD94; log2FC = 1.50, p_adj_ = 1.9 × 10^-12^), KLRG1 (log2FC = 1.35, p_adj_ = 4.4 × 10^-7^), NKG7 (log2FC = 0.75, p_adj_ = 4.1 × 10^-6^), FCRL6 (log2FC = 1.89, p_adj_ = 2.1 × 10^-10^), and CCL5 (log2FC = 0.57, p_adj_ = 5.7 × 10^-8^), while low-shift clones showed higher expression of IL7R, ICOS, and NELL2. This pattern persisted at day 119, with high-shift clones sustaining KLRD1, KLRG1, and KLRK1 (NKG2D; log2FC = 1.08, p_adj_ = 0.002), and low-shift clones showing higher ICOS, IL7R, LAIR1, and interferon-response genes (IFITM2, IFI27). In DFCI2 CD8+ effector memory T cells and DFCI1 stress-signature T cells, no FDR-significant transcriptional programs were identified, indicating that the remodeling signal in those compartments, while detectable at the clonotype level, does not correspond to a discrete transcriptional effector state.

**Figure S4.**
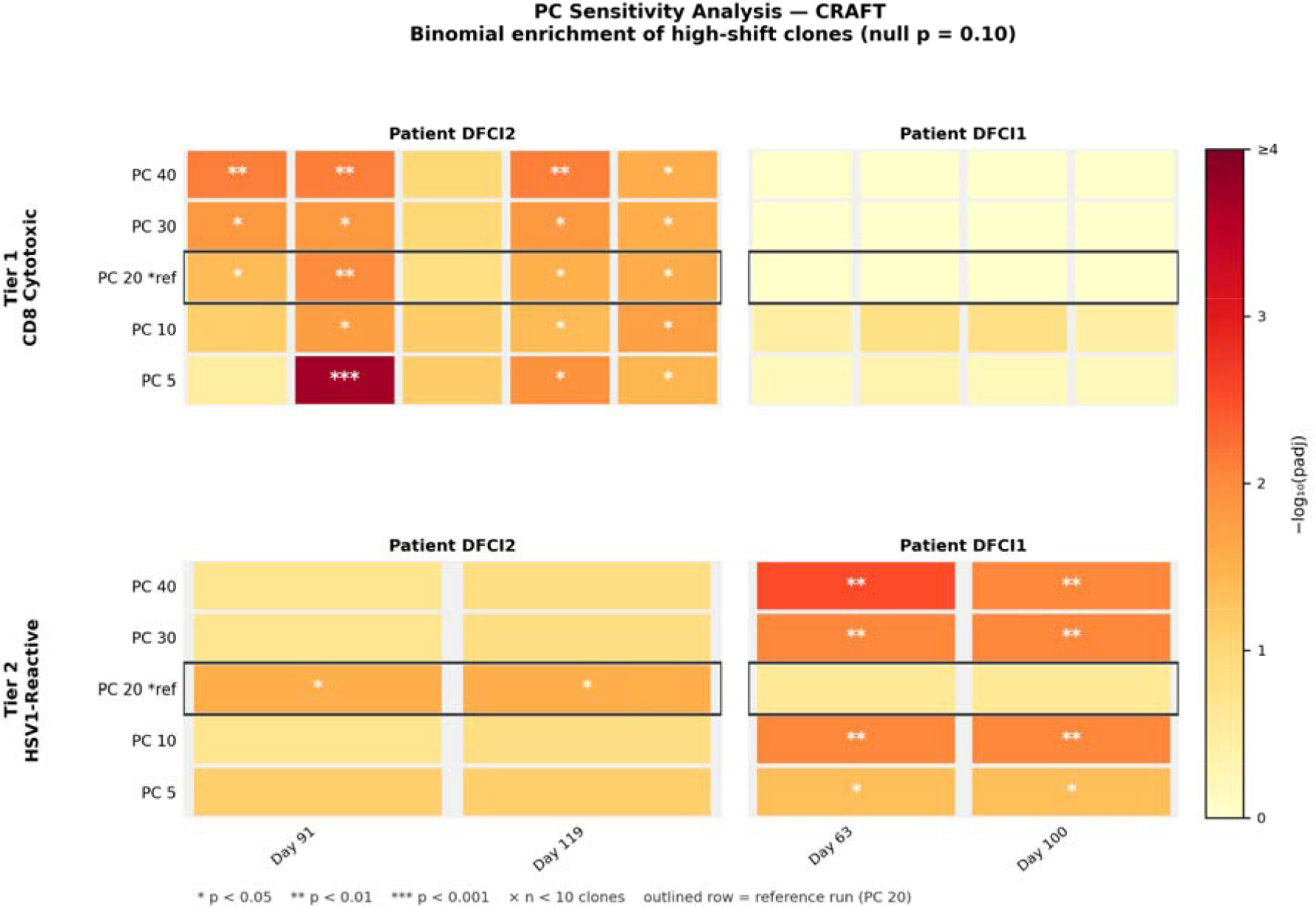
Sensitivity of tail fraction distribution in the GBM dataset for 5,10, 20, 30 and 40 PCs

**Figure S5.**
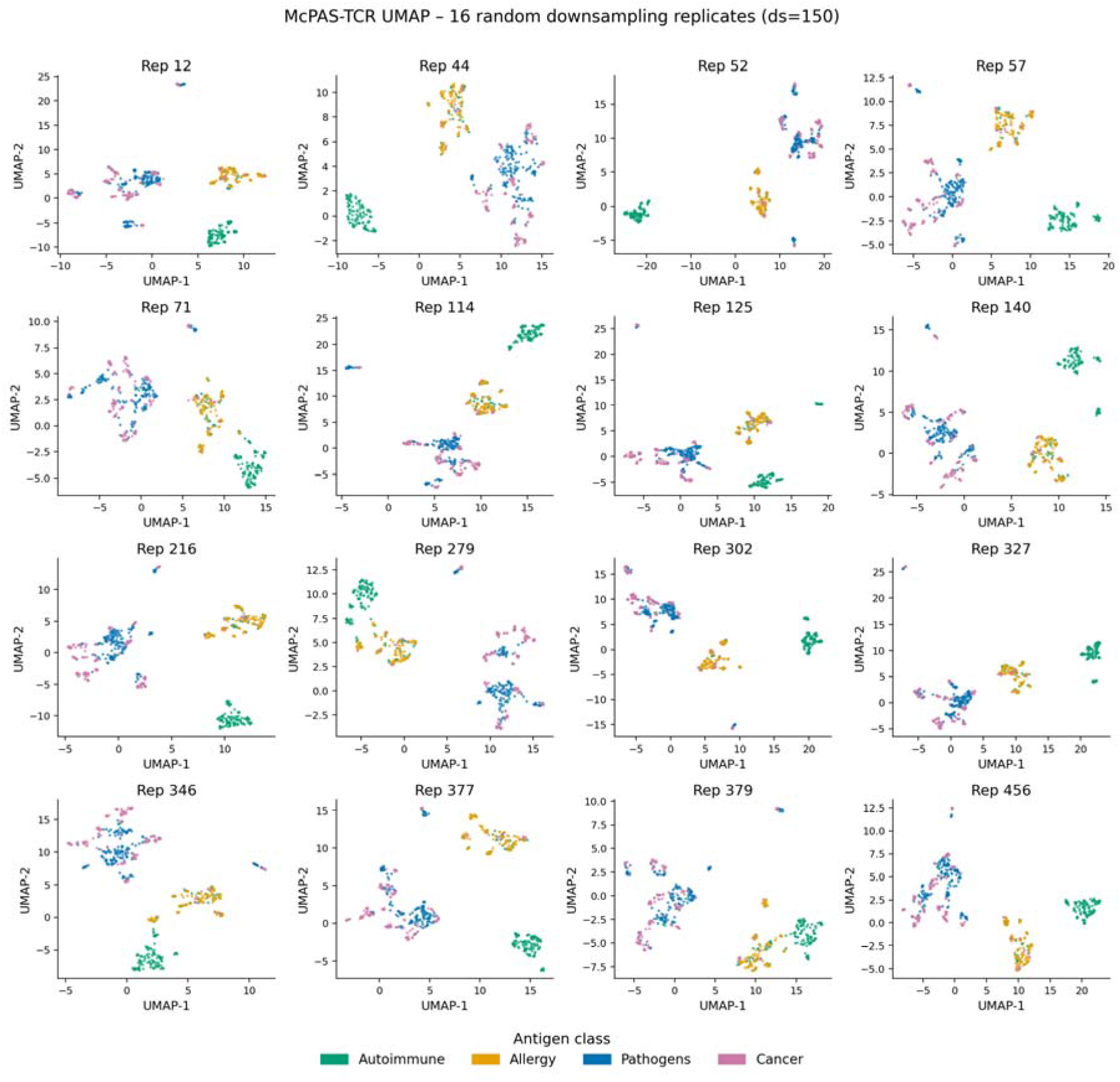
Antigen clustering across downsampled replicates in the McPAS-TCR dataset. UMAP visualizations of CRAFT embeddings for representative balanced subsamples (150 sequences per antigen class) and for the full unbalanced dataset (n = 5,279), colored by antigen class (Pathogens, Autoimmune, Cancer, Allergy). Across replicates, autoimmune- and pathogen-specific TCRs consistently form distinct, compact clusters, with cancer-associated TCRs positioned peripherally to the pathogen cluster. A third cluster comprises a mixture of allergy- and autoimmune-related TCRs. The consistency of cluster structure across independent downsampled replicates confirms that the observed antigen-class organization is robust to sampling variability and not driven by individual sequences or class-size imbalance.

**Table S1.** Model training hyperparameters across curriculum phases. CRAFT training used a multi-phase curriculum strategy that progressively increased sequence complexity. For each phase, the table reports dropout rate, learning rate, warmup steps, total steps, weight decay, optimizer decay, and batch size. Transitions between phases were triggered after convergence criteria were met on the validation loss.

| Phase | Dropout | Learning Rate | Warmup Steps | Total Steps | Weight Decay | Optimizer Decay | Batch Size |
| --- | --- | --- | --- | --- | --- | --- | --- |
| Phase 1 | 0.05 | $2e^{-4}$ | 10000 | 105000 | 0.09 | 0.10 | 1024 |
| Phase 2 (Epoch 1) | 0.10 | $1e^{-4}$ | 9000 | 105000 | 0.09 | 0.05 | 1024 |
| Phase 2 (Epoch 2) | 0.10 | $1e^{-4}$ | 3000 | 105000 | 0.09 | 0.05 | 1024 |
| Phase 2 (Epoch 3) | 0.10 | $5e^{-5}$ | 3000 | 105000 | 0.09 | 0.05 | 1024 |
| Phase 3 (Epoch 1) | 0.12 | $5e^{-5}$ | 10000 | 105000 | 0.09 | 0.01 | 1024 |
| Phase 3 (Epoch 2) | 0.13 | $5e^{-5}$ | 3000 | 105000 | 0.09 | 0.005 | 1024 |

**Table S2.** Distribution of treatment arms across pathological response groups in the OCSCC cohort. Patients received neoadjuvant anti-PD-1 (nivolumab) monotherapy or combination anti-PD-1 plus a single dose of anti-CTLA-4 (ipilimumab). Response groups were defined by pathological response: High (50-100% pathological response), Medium (10-49%), Low (0-9%), per Luoma et al. (2022). Treatment arms were not balanced across response groups: 4 of 6 high responders received combination therapy, while 4 of 5 low responders received monotherapy. Tables are provided for both the full Luoma cohort (28 evaluable patients) and the CRAFT analysis subset (18 patients with complete PBMC sampling at all three timepoints).

**For the full Luoma cohort (28 evaluable patients):**
|  | High | Medium | Low | Total |
| --- | --- | --- | --- | --- |
| Monotherapy (Nivo) | 2 | 5 | 6 | 13 |
| Combination (Nivo + Ipi) | 5 | 6 | 4 | 15 |
| Total | 7 | 11 | 10 | 28 |

**For the CRAFT analysis subset (18 patients with complete PBMC at B1, B2, B3):**
|  | High | Medium | Low | Total |
| --- | --- | --- | --- | --- |
| Monotherapy (Nivo) | 2 | 4 | 4 | 10 |
| Combination (Nivo + Ipi) | 4 | 3 | 1 | 8 |
| Total | 6 | 7 | 5 | 18 |

**Table S3.**
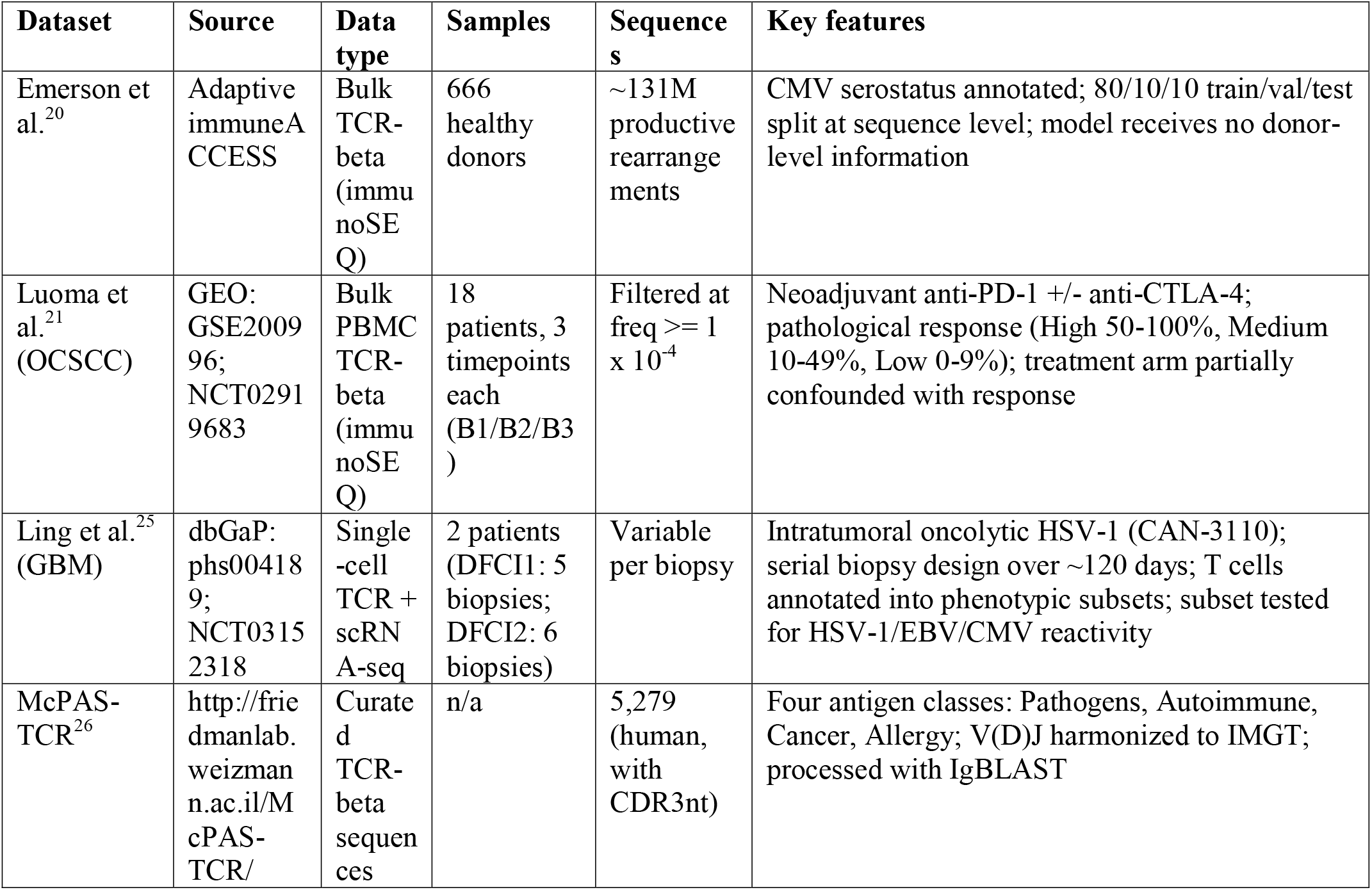
Detailed dataset descriptions.

