## Supplementary_Information for "A generative reference grammar of healthy TCR repertoires reveals cancer-associated immune remodeling"

**Compartment-specific longitudinal immune remodeling: extended analysis**

**Reactivity-stratified displacement in DFCI2.** To assess whether displacement in the clinically stable patient (DFCI2) reflected antigen-specific clonal dynamics, we stratified CD8+ cytotoxic clonotypes by TCR reactivity. HSV-1-reactive clonotypes showed progressive broadening of distance distributions from day 30 onward, with a distinct subset of clonotypes exceeding the baseline 90th-percentile threshold. EBV-reactive and non-reactive clonotypes remained near baseline, consistent with antigen-specific clonal expansion selectively reshaping the repertoire in response to the oncolytic virus rather than reflecting a global inflammatory perturbation. Restricting the binomial tail enrichment test to HSV-1-reactive clonotypes alone, significant enrichment was observed at days 91 and 119 (p_adj = 0.029; tail fractions 30% and 38%), confirming that displacement signal persists within this antigen-specific subpopulation (Figure 3a,b)

**Differential gene expression analysis.** To determine whether geometric displacement in CRAFT embedding space corresponds to transcriptional changes, we compared gene expression between highly displaced (above the 75th percentile of baseline distance) and minimally displaced (below the 25th percentile) CD8+ cytotoxic clonotypes in DFCI2 at days 91 and 119 using Wilcoxon rank-sum tests on log-normalized counts. Highly displaced clonotypes were enriched for markers of terminal effector differentiation: KLRD1 (CD94; log2FC = 1.50), KLRG1 (log2FC = 1.35), NKG7, FCRL6, and CCL5. Low-displacement clonotypes expressed memory-associated markers including IL7R and ICOS. This pattern persisted at day 119 (Fig. 3d in the current manuscript; Extended Data Fig. 6 in the revised manuscript).

The upregulation of KLRD1 and KLRG1, canonical markers of terminally differentiated cytotoxic effectors, together with the NK-like cytotoxicity program (NKG7, FCRL6) indicates that CRAFT displacement captures a biologically meaningful transition from quiescent memory toward active antiviral effector function. That low-displacement clonotypes retain IL7R and ICOS expression is consistent with their maintenance in a memory or progenitor state, further supporting the interpretation that displacement in embedding space tracks effector differentiation. We confirmed a similar pattern when comparing clones above the 90th percentile from baseline versus clones below the 10th percentile (Figure S3).

**Stress-signature T cell dynamics in DFCI1.** The progressing patient (DFCI1) had no radiological response, although he did have a pathological response/treatment effect. In this patient, neither CD8+ compartment showed systematic displacement. However, the stress-signature compartment exhibited a strikingly different trajectory: large clonal expansion at day 17 with sustained enrichment of displaced clonotypes through day 63 (p_adj = 7.6 x 10^-31, 1.6 x 10^-4, 2.8 x 10^-1). This pattern is consistent with progressive T cell dysfunction under persistent tumor pressure, where stress-associated T cells expand but fail to mount an effective cytotoxic response, a hallmark of the exhausted tumor microenvironment in glioblastoma. (Figure 3a,b).
